# NRF2 co-opts the SWI/SNF complex to drive liver cell plasticity

**DOI:** 10.64898/2026.02.09.704724

**Authors:** Athena Jessica S Ong, Chapman CH Wong, Anthony P Karamalakis, Kimberley J Evason, Kristin K Brown, Andrew G Cox

**Author notes:** Equal contribution (Kristin K Brown, Andrew G Cox).

## Abstract

Nuclear factor erythroid 2-related factor 2 (NRF2) is a transcription factor that plays a role in the regulation of redox homeostasis and cellular metabolism. Activating mutations in the NRF2 pathway have been identified in approximately 15% of liver cancer patients. However, the mechanisms by which NRF2 promotes liver tumorigenesis are poorly understood. Employing a transgenic zebrafish model with hepatocyte-specific, inducible expression of a clinically relevant constitutively active NRF2 mutant (NRF2^T80K^), we show that constitutive activation of NRF2 drives hepatocyte to cholangiocyte transdifferentiation. Importantly, we demonstrate that NRF2 affects liver cell plasticity in a cell-autonomous, evolutionarily conserved, and reversible manner. Utilizing an epigenetic-focused chemical screen, the BRG1/BRM inhibitor FHD-286 was identified as a potent suppressor of NRF2-driven transdifferentiation. Overall, our study reveals a novel role for NRF2 in the regulation of liver cell plasticity during tumour initiation and identifies a therapeutic approach to overcome the oncogenic activity of NRF2.

## Introduction

Liver cancer is the third deadliest cancer worldwide and has a dismal 5-year survival rate of approximately 18%^1^. The main types of liver cancer are hepatocellular carcinoma (HCC; 75%-85%) and cholangiocarcinoma (CCA; 10%-15%)^1^. While HCC originates from hepatocytes, CCA can originate from cholangiocytes, hepatocytes, or other cell types in the liver^2–5^. Lineage tracing studies have shown that CCA, induced via carcinogen exposure or oncogenic pathway activation, can originate from hepatocytes that have transdifferentiated to a biliary cell fate^3,6–9^. The emerging consensus suggests that the transdifferentiation of hepatocytes to cholangiocytes constitutes a critical initial step in the formation of CCA.

Analysis of sequencing data from the International Cancer Genome Consortium (ICGC) and The Cancer Genome Atlas (TCGA), has revealed that the NRF2 pathway is genetically altered in up to 15% of HCC cases^10,11^. NRF2 is a transcription factor sensitive to a variety of environmental stressors, in particular oxidative stress^12–14^. Therefore, the incidence of NRF2 pathway activation in liver cancer is likely significantly higher given that oxidative stress is a characteristic feature of pre-malignant chronic liver diseases, including metabolic dysfunction-associated steatotic liver disease (MASLD), which are impacting an increasing percentage of the global population^15,16^. Consistent with this, the expression of a NRF2 target gene signature is more pervasive in TCGA data than would be expected based on the incidence of pathway mutations in HCC^17^. An increasing body of experimental evidence supports an important role for NRF2 pathway activation in the formation of liver cancer, although the mechanisms underpinning NRF2-driven tumorigenesis are poorly understood^15,18–20^.

Clinically relevant hotspot mutations in NRF2 are present in the DLG or ETGE motifs within the NRF2-ECH homology 2 (Neh2) domain of NRF2, which serve as evolutionarily conserved binding sites for Kelch-like ECH-associated protein 1 (KEAP1), a negative regulator of NRF2^21–23^. Mutations within these motifs disrupt binding with KEAP1 leading to the constitutive activation of NRF2^14^. For example, a particularly prevalent NRF2 mutation, NRF2^T80K^, gives rise to an EKGE motif that no longer binds KEAP1.

In this study, we have developed a transgenic zebrafish model with hepatocyte-specific, inducible expression of NRF2^T80K^ to investigate the effect of NRF2 activation on liver tumour initiation. We show that NRF2 expression promotes expansion of the cholangiocyte compartment and development of a glandular phenotype reminiscent of CCA. We find that cholangiocyte expansion results from a cell-autonomous and evolutionarily conserved mechanism of hepatocyte to cholangiocyte transdifferentiation. Mechanistically, we reveal that NRF2-driven cell plasticity is dependent on the SWI/SNF complex and represents a vulnerability that can be therapeutically targeted to inhibit NRF2-driven liver tumour initiation.

## Results

### Constitutive NRF2 activation promotes liver cell plasticity *in vitro* and *in vivo*

To investigate the role of constitutively active NRF2 in liver cancer, we first examined our previously published RNA sequencing (RNA-Seq) dataset (GSE230611) comparing wild-type and KEAP1-deficient HepG2 liver cancer cells^24^. Interestingly, gene set enrichment analysis (GSEA) revealed differential expression of pathways related to liver cell fate. Specifically, gene sets associated with HNF4A, a transcription factor regulating hepatocyte maturation and function^25–28^, were downregulated in the context of KEAP1 deficiency (Fig. 1A). In contrast, gene sets related to the acquisition of cholangiocyte fate were upregulated (Fig. 1A). Downregulation of HNF4A was confirmed at the level of protein expression (Fig. 1B). To determine whether changes in liver cell fate were an early response to NRF2 activation, HepG2 cells were engineered to enable doxycycline (DOX)-inducible expression of the oncogenic NRF2^T80K^ mutant (TO:NRF2^T80K^). DOX dosage was optimised to achieve comparable levels of NRF2 activation between the two cell line models (Fig. S1A,B). As expected, genes associated with NRF2 pathway activation were upregulated in DOX-treated TO:NRF2^T80K^ cells (Fig. 1C,D). Importantly, downregulation of hepatocyte-associated genes, reduced expression of HNF4A, and upregulation of cholangiocyte-associated genes was observed in DOX-treated TO:NRF2^T80K^ cells (Fig. 1D-F), concordant with results from KEAP1-deficient cells. Together, these data suggest that constitutive activation of NRF2 promotes liver cell plasticity leading to loss of HNF4A and acquisition of a cholangiocyte-like fate.

**Figure 1.**
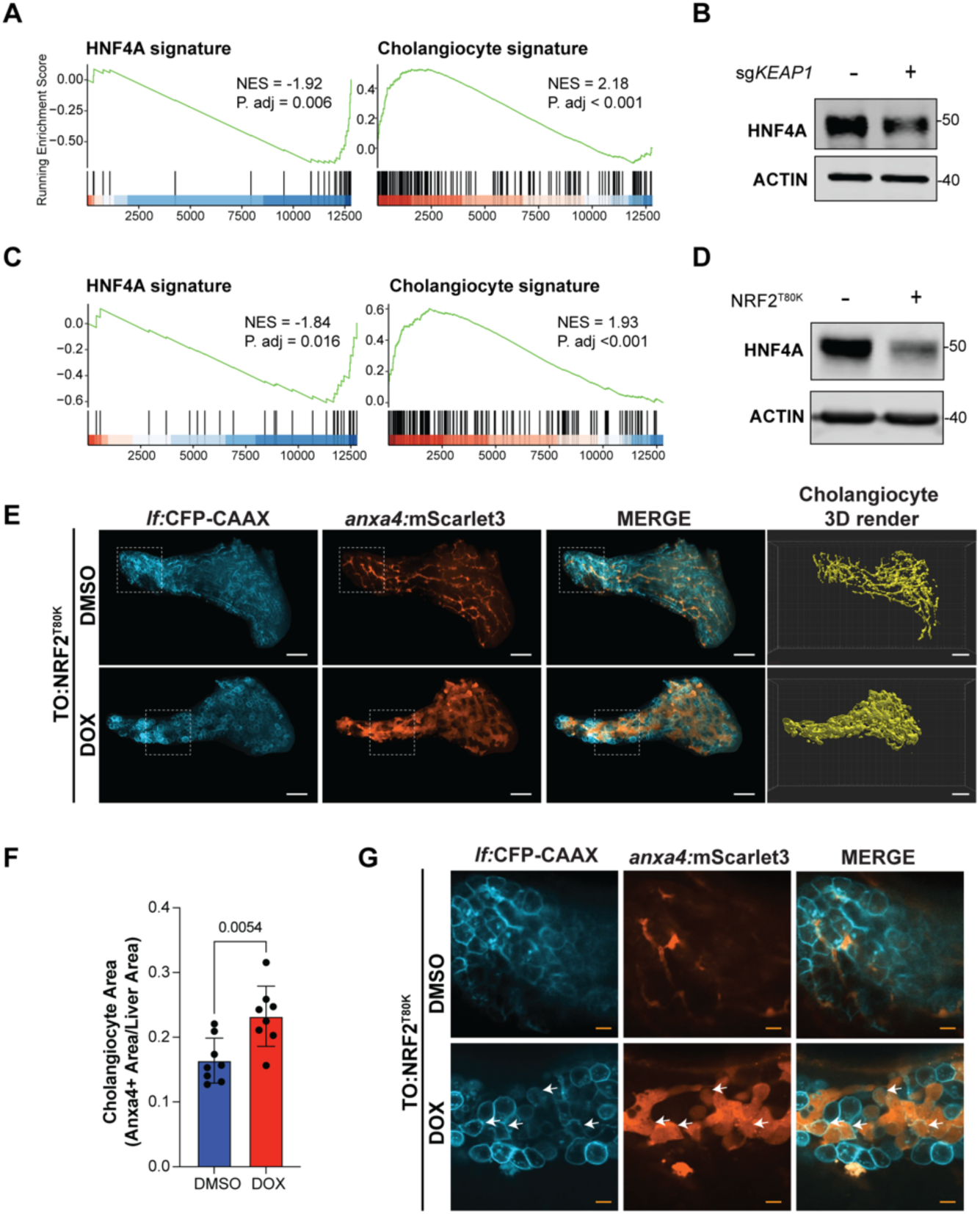
Constitutive NRF2 activation promotes liver cell plasticity *in vitro* and *in vivo*. (A) Gene set enrichment analysis (GSEA) plots derived from RNA-Seq analysis of Cas9-expressing HepG2 cells transduced with sgRNA targeting *KEAP1* (sg*KEAP1*) versus cells transduced with sgRNA targeting the safe harbour locus AAVS1 (sgAAVS1), n=4. HNF4A signature, Gene Set SUMI_HNF4A_TARGETS; Cholangiocyte signature, Gene set AIZARANI_LIVER_C39_EPCAM_POS_BILE_DUCT_CELLS_4. (B) Representative immunoblot analysis of sgAAVS1 and sg*KEAP1* HepG2 cells. (C) GSEA plot derived from RNA-Seq analysis of HepG2 cells harbouring inducible expression of EGFP (Control) versus cells harbouring inducible expression of NRF2^T80K^ (TO:NRF2^T80K^) after 4 days of DOX-treatment, n=3. NRF2 pathway signature corresponds to Gene Set WP_NRF2_PATHWAY. (D) Volcano plot of differentially expressed genes (DEGs) identified by RNA-Seq analysis of TO:NRF2^T80K^ versus Control HepG2 cells after 4 days of DOX treatment, n=3. Significant DEGs are highlighted in pink. Select canonical NRF2 target genes are highlighted in blue. Cholangiocyte markers are highlighted in green. (E) GSEA plots derived from RNA-Seq analysis of Control versus TO:NRF2^T80K^ HepG2 cells, n=3. HNF4A signature, Gene Set SUMI_HNF4A_TARGETS; Cholangiocyte signature, Gene set AIZARANI_LIVER_C39_EPCAM_POS_BILE_DUCT_CELLS_4. (F) Representative immunoblot analysis of Control and TO:NRF2^T80K^ HepG2 cells after 4 days of DOX treatment. (G) Representative confocal images of hepatocyte membrane (*lf*:CFP-CAAX, cyan), cholangiocytes (*anxa4*:mScarlet3, red), and a 3D-rendered mask of the cholangiocyte network derived from TO:NRF2^T80K^;*anxa4*:mScarlet3 zebrafish larvae treated with DMSO or DOX from 5 to 10 days post-fertilisation (dpf). White scale bars represent 50 µm. (H) Quantification of mScarlet3 (cholangiocyte) area normalized to CFP (liver) area, n=8. (I) Magnification of liver sections denoted by hashed lines in (G). White arrows indicate cells expressing hepatocyte (cyan) and cholangiocyte (red) markers. Scale bars represent 10 µm.

To determine whether NRF2 influences cell fate *in vivo*, a zebrafish model with hepatocyte-specific, DOX-inducible expression of NRF2^T80K^ was generated. TO:NRF2^T80K^ zebrafish express a membrane-bound form of CFP that marks hepatocytes. To validate the model, TO:NRF2^T80K^ zebrafish were crossed to a NRF2 reporter line (*gstp1*:EGFP) and hepatocyte-specific NRF2 activation was exclusively observed in DOX-treated larvae (Fig. S1C). To enable real-time visualization of cholangiocyte fate *in vivo*, a cholangiocyte reporter was generated by knocking-in a cleavable mScarlet3-p2a at the transcription start site of the cholangiocyte marker annexin A4 (*anxa4*; Fig. S1D). Given that the zebrafish liver is fully functional by 5 days post-fertilisation (dpf)^29^, the effects of NRF2 activation were examined in compound TO:NRF2^T80K^;*anxa4*:mScarlet3 larvae treated with DOX from 5 to 10 dpf. Mirroring our *in vitro* observations, a dramatic increase in cholangiocyte density was observed in the context of NRF2 activation (Fig. 1G,H). Interestingly, cholangiocytes in DOX-treated zebrafish exhibited an atypical cuboidal morphology that was distinct from the more classical tubular morphology observed in the control setting (Fig. 1G,I). Moreover, multiple hepatocytes also expressed *anxa4* in DOX-treated larvae (Fig. 1I). Importantly, DOX treatment did not impact the hepatocyte or cholangiocyte compartments in wild-type *anxa4*:mScarlet3 larvae (Fig. S1E). These observations were phenocopied in *keap1a*/*keap1b* crispants (Fig. S1F). Collectively, these results suggest that hepatocyte-specific activation of NRF2 promotes an expansion of the cholangiocyte compartment *in vivo*.

### NRF2 promotes transdifferentiation of hepatocytes to cholangiocytes

Given that cell fate plasticity is an inherent feature of organogenesis, we turned to investigate the impact of NRF2 activation in fully developed, adult TO:NRF2^T80K^ zebrafish. Hematoxylin and Eosin (H&E) staining of histological sections from adult TO:NRF2^T80K^ zebrafish exposed to DOX for 4 weeks showed a dramatic increase in basophilic staining of the hepatic parenchyma and appearance of glandular structures, reminiscent of features associated with CCA (Fig. 2A). Moreover, immunofluorescent staining revealed an expansion of the cholangiocyte compartment and altered cholangiocyte morphology in TO:NRF2^T80K^ zebrafish following DOX-treatment (Fig. 2A). Furthermore, bilineage cells that were positive for both hepatocyte and cholangiocyte markers were observed in TO:NRF2^T80K^ livers (Fig. 2B). In alignment with *in vitro* data, bulk RNA-Seq analysis revealed that NRF2 activation induces expression of bona fide NRF2 target genes, downregulation of hepatocyte genes and upregulation of cholangiocyte genes (Fig. 2C and Fig. S2A). To examine the consequences of NRF2 activation at the single cell level, single-cell RNA-Seq (scRNA-Seq) was performed. As anticipated, Uniform Manifold Approximation and Projection (UMAP) visualization confirmed expression of the NRF2 target gene *glutathione S-transferase pi 1* (*gstp1*) exclusively in the DOX-treated group (Fig. S2B). Integrated clustering of DMSO- and DOX-treated groups identified 16 transcriptionally distinct clusters (Fig. 2D,E). Specific cell types were assigned based on the expression of previously identified cell type markers (Supplementary Table 1)^30^. Quantification of the relative abundance of the major liver cell types revealed that NRF2 activation promotes a loss of hepatocytes, a gain of cholangiocytes, and the appearance of a novel cluster with mixed hepatocyte and cholangiocyte features, consistent with the phenotype observed by immunofluorescent staining (Fig. 2F,G). Interestingly, clusters with mixed hepatocyte and cholangiocyte markers in the context of NRF2 activation bridged the hepatocyte and cholangiocyte clusters, suggesting the emergence of bilineage hepatocytes acquiring cholangiocyte features (Fig. 2G). Ductular reaction (DR) is a term used to describe hyperplasia of cholangiocytes, which is a common reaction to liver injury and associated with induction of apoptosis, inflammatory cell infiltration, and fibrosis^31–34^. The absence of fibrosis (Fig. S2C) and apoptosis (Fig. S2D) in livers from DOX-treated TO:NRF2^T80K^ zebrafish suggests that DR is not the cause of cholangiocyte expansion.

**Figure 2.**
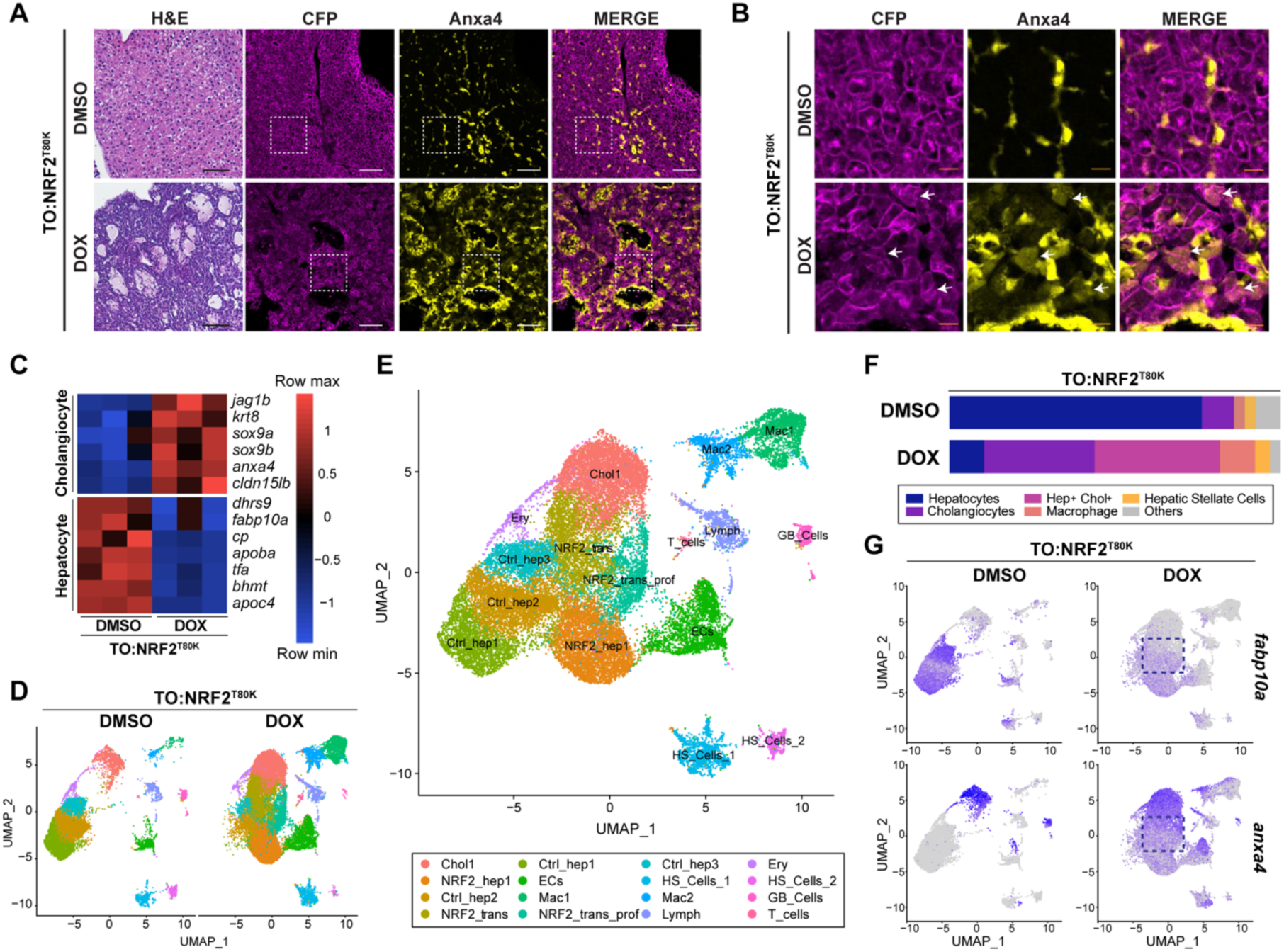
NRF2 promotes transdifferentiation of hepatocytes to cholangiocytes. (A) Representative H&E and immunofluorescent staining of CFP (hepatocyte, magenta) and Anxa4 (cholangiocyte, yellow) in liver sections from adult TO:NRF2^T80K^ zebrafish after 4 weeks treatment with DMSO or DOX. Scale bars represent 50 µm. (B) Magnification of liver sections denoted by hashed lines in (A). White arrows indicate cells that express CFP (hepatocyte, magenta) and Anxa4 (cholangiocyte, yellow) markers. Scale bars represent 10 µm. (C) Heatmap of hepatocyte and cholangiocyte gene expression in livers from adult TO:NRF2^T80K^ zebrafish after 4 weeks treatment with DMSO or DOX as determined by bulk RNA-Seq analysis, n=3. (D) Uniform Manifold Approximation and Projection (UMAP) visualization of scRNA-Seq data obtained from livers from adult TO:NRF2^T80K^ zebrafish after 3 weeks treatment with DMSO or DOX. (E) UMAP visualization from the joint clustering of scRNA-Seq data obtained from livers from adult TO:NRF2^T80K^ zebrafish after 3 weeks treatment with DMSO or DOX. Chol: cholangiocytes; ECs: Endothelial cells; Ery: Erythrocytes; Hep: Hepatocytes; HS_Cells: Hepatic stellate cells; Lymph: Lymphocytes; Mac: Macrophages; Nrf2_trans: Transdifferentiating cells; GB_Cells: Gall bladder cells. (F) Stacked bar chart showing the relative proportions of the major cell types in livers from adult TO:NRF2^T80K^ zebrafish treated with DMSO or DOX for 3 weeks. (G) UMAP visualization of *fabp10a* (hepatocyte) and *anxa4* (cholangiocyte) expression in livers from adult TO:NRF2^T80K^ zebrafish treated with DMSO or DOX for 3 weeks. Hashed boxes outline cells expressing both markers.

### NRF2 is required to maintain altered liver cell fate identity

To determine the reversibility of NRF2-driven transdifferentiation DOX pulse-chase experiments were performed in adult TO:NRF2^T80K^ zebrafish (Fig. 3A). Zebrafish were exposed to DOX or vehicle (DMSO) for 7 or 14 days, at which point gland-like structures were present and there was an evident expansion of the cholangiocyte compartment (Fig. 3B,C). This was followed by a washout period of 7 or 28 days. H&E staining revealed a reversion of NRF2^T80K^-transformed livers to a histologically normal state 28 days after DOX removal (Fig. 3B,C). Moreover, immunofluorescent staining highlighted an indistinguishable difference in hepatocyte and cholangiocyte compartments between control zebrafish and NRF2-transformed livers 28 days after DOX removal (Fig. 3C). Periodic Acid-Schiff (PAS) staining, which detects glycogen storage in mature hepatocytes, confirmed loss of hepatocyte function following DOX exposure and subsequent recovery of hepatocyte function 28 days after DOX removal (Fig. S3A). Bulk RNA-Seq analysis demonstrated that, by the end of the 28 day washout period, the liver transcriptome had nearly completely reverted to the homeostatic state (Fig. 3D). This included a normalisation in the expression of NRF2 target genes, hepatocyte-associated genes and cholangiocyte-associated genes (Fig. 3E,F and Fig. S3B). The reversibility of NRF2-induced changes in liver cell fate were confirmed in pulse-chase studies employing TO:NRF2^T80K^ HepG2 cells (Fig. 3G and Fig. S3C). Together, these data demonstrate that NRF2 is required to maintain altered liver cell fate identity in a cell autonomous and evolutionarily conserved manner.

**Figure 3.**
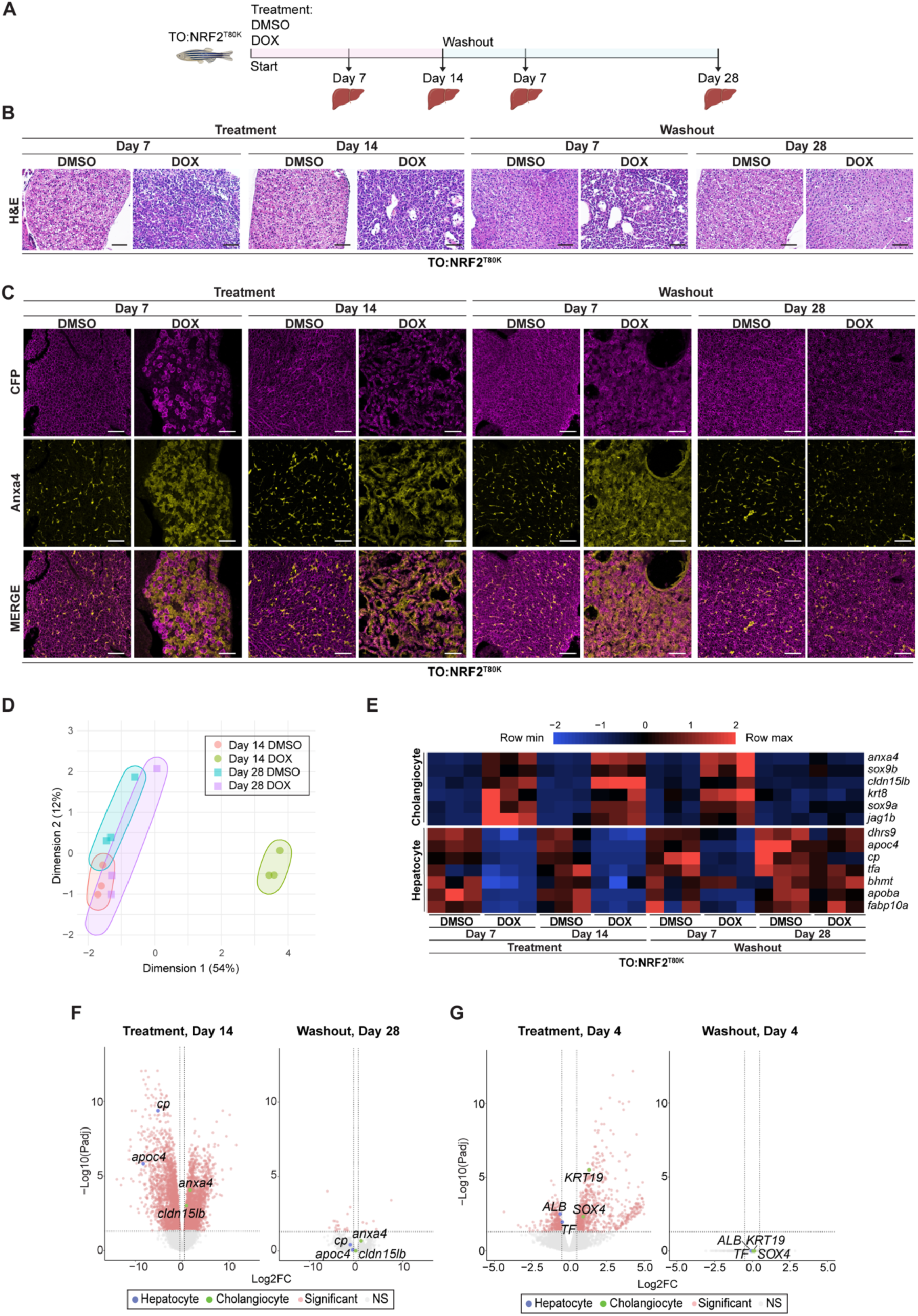
NRF2 is required to maintain altered liver cell fate identity. (A) Schematic of DMSO/DOX pulse-chase experiments in adult TO:NRF2^T80K^ zebrafish. (B) Representative H&E-staining in liver sections from adult TO:NRF2^T80K^ zebrafish after 7- or 14-days treatment with either DMSO or DOX, followed by 7- or 28-days withdrawal of DMSO or DOX. Scale bars represent 50 µm. (C) Representative immunofluorescent staining of CFP (hepatocyte, magenta) and Anxa4 (cholangiocyte, yellow) in liver sections from adult TO:NRF2^T80K^ zebrafish after 7- or 14-days treatment with either DMSO or DOX, followed by 7- or 28-days withdrawal of DMSO or DOX. Scale bars represent 50 µm. (D) Multidimensional scaling (MDS) plot of bulk RNA-Seq data obtained from livers isolated from adult TO:NRF2^T80K^ zebrafish after 14 days treatment with either DMSO or DOX, followed by 28-days withdrawal of DMSO or DOX, n=3. (E) Heatmap of hepatocyte and cholangiocyte gene expression in livers isolated from adult TO:NRF2^T80K^ zebrafish after 7- or 14-days treatment with either DMSO or DOX, followed by 7- or 28-days withdrawal of DMSO or DOX, as determined by bulk RNA-Seq analysis, n=3. (F) Volcano plot of DEGs identified by bulk RNA-Seq analysis of livers isolated from adult TO:NRF2^T80K^ zebrafish treated with DOX versus DMSO for 14 days (left), followed by withdrawal of DOX versus DMSO for 28 days (right), n=3. Significant DEGs are highlighted in pink. Hepatocyte markers are highlighted in blue. Cholangiocyte markers are highlighted in green. (G) Volcano plot of DEGs identified by RNA-Seq analysis of TO:NRF2^T80K^ versus Control HepG2 cells after 4 days of DOX treatment (left), followed by withdrawal of DOX for 4 days (right). Significant DEGs are highlighted in pink. Hepatocyte markers are highlighted in blue. Cholangiocyte markers are highlighted in green.

### An epigenetic-focused inhibitor screen reveals that NRF2-driven liver cell plasticity is mediated by BRG1/BRM

Cell fate plasticity is recognised to be dependent on epigenetic remodelling^35–38^. To identify candidate epigenetic regulators of NRF2-driven liver cell plasticity, a chemical library comprising 38 clinically relevant small molecule inhibitors of key epigenetic regulators was employed in an *in vivo* screen (Fig. S4A, Supplementary Table 2). Briefly, TO:NRF2^T80K^; *anxa4*:mScarlet3 transgenic larvae were exposed to DMSO or DOX in the presence or absence of the epigenetic inhibitors for 4 days before confocal analysis of liver cell fate. One of the most potent suppressors of NRF2-driven transdifferentiation was the BRG1/BRM inhibitor FHD-286 (Fig. 4A)^39^. BRG1 and BRM constitute the two catalytic subunits of SWI/SNF chromatin remodelling complexes, which have been shown to play a role in the maintenance of cell fate^40,41^. Interestingly, biochemical studies have previously suggested that NRF2 interacts with BRG1^42,43^. Suppression of NRF2-driven transdifferentiation by FHD-286 in the larval setting was confirmed in a validation cohort. Importantly, treatment of a DMSO-control group treated with FHD-286 confirmed that inhibition of BRG1/BRM was not associated with signs of cholangiotoxicity, which could be interpreted as suppression of transdifferentiation (Fig. 4B,C). To overcome potential complications associated with inhibition of SWI/SNF complexes during organogenesis, the impact of FHD-286 on NRF2-driven changes in cell fate plasticity were also examined in adult zebrafish. Consistent with larval studies, FHD-286 effectively suppressed NRF2-driven changes in liver architecture and expansion of the cholangiocyte compartment (Fig. 4D,E). Importantly, while BRG1/BRM inhibition altered the transcriptional landscape in DOX-treated TO:NRF2^T80K^ zebrafish, minimal changes in gene expression were observed in the absence of NRF2 activation (Fig. 4F). Interestingly, while able to block NRF2-driven changes in expression of genes related to liver cell fate, FHD-286 did not completely suppress NRF2 target gene expression (Fig. S4B,C). Finally, analysis of TCGA Cancer Dependency Map (TCGA_DEPMAP_)^44^ revealed an essential role for *SMARCA4* (gene encoding BRG1) and *SMARCA2* (gene encoding BRM) in the context of *KEAP1* loss-of-function (Fig. 4G). Together, these data highlight a role for SWI/SNF-dependent chromatin remodelling complex in regulating liver cell plasticity in the context of NRF2 pathway activation.

**Figure 4.**
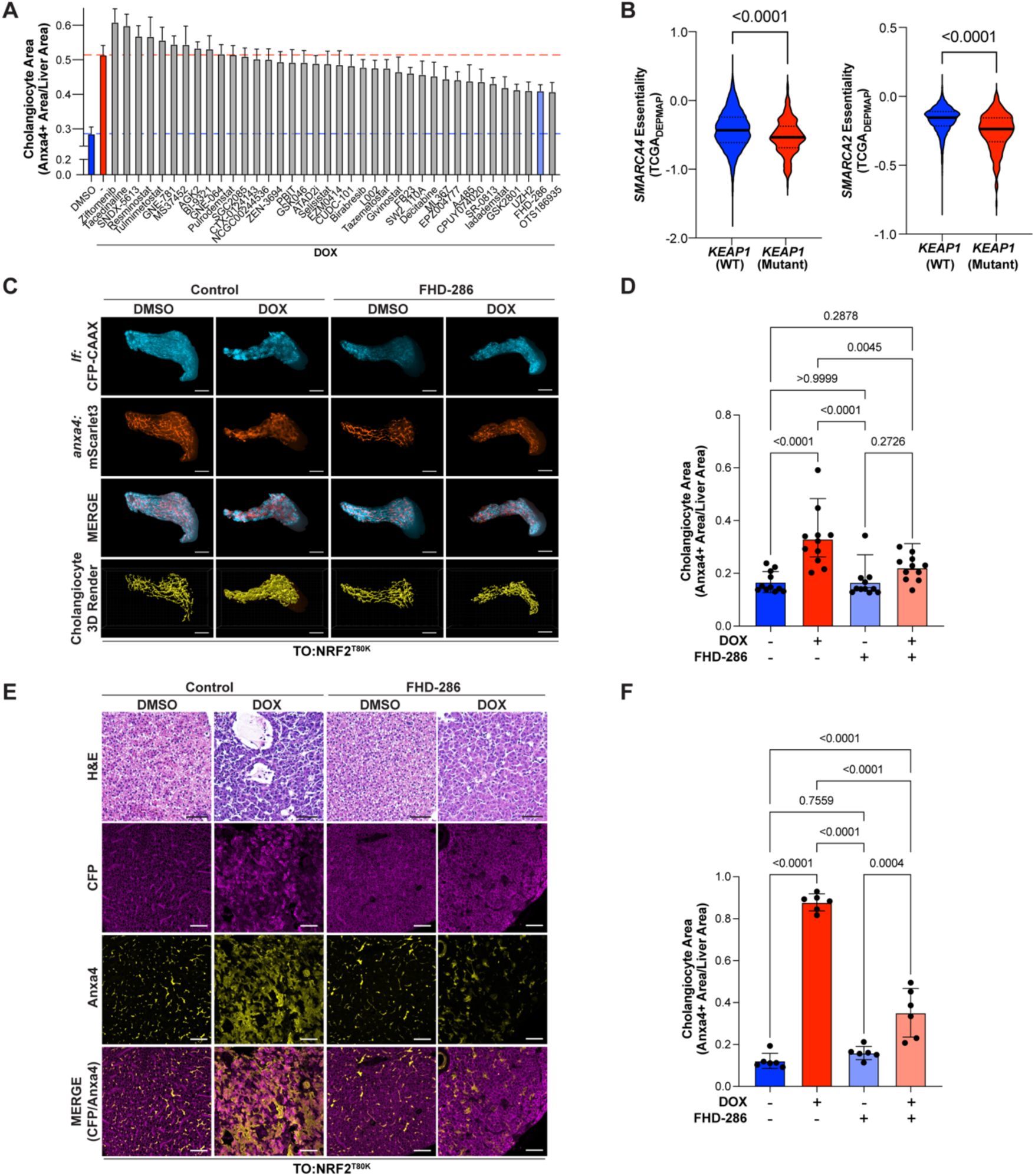
An epigenetic-focused inhibitor screen reveals that NRF2-driven liver cell plasticity is mediated by BRG1/BRM. (A) Quantification of cholangiocyte area (mScarlet3), as a ratio of liver area (CFP) area, from an *in vivo* chemical suppressor screen using an epigenetic-focused library of small molecule inhibitors. The blue dashed line indicates the mean of the DMSO-treated group, whereas the red dashed line indicates the mean of the DOX-treated group. (B) Representative confocal images of CFP (hepatocyte, cyan), mScarlet3 (cholangiocyte, red), and a 3D-rendered mask of the cholangiocyte network from TO:NRF2^T80K^;*anxa4*:mScarlet3 zebrafish larvae with DMSO or DOX in the absence or presence of 100 nM FHD-286 from 5 to 10 dpf. White scale bars represent 50 µm. (C) Quantification of mScarlet3 (cholangiocyte) area normalized to CFP (liver) area, n=11. (D) Representative H&E and immunofluorescent staining of CFP (hepatocyte, magenta) and Anxa4 (cholangiocyte, yellow) in liver sections from adult TO:NRF2^T80K^ zebrafish treated with DMSO or DOX in the absence or presence of 250 nM FHD-286 for 14 days. (E) Quantification of Anxa4 (cholangiocyte) area normalized to CFP (liver) area, n=6. (F) Volcano plot of DEGs identified by bulk RNA-Seq analysis of livers isolated from adult TO:NRF2^T80K^ zebrafish treated with DOX (left) or DMSO (right) in the presence versus absence of 250 nM FHD-286 for 14 days. Significant DEGs are highlighted in pink. Hepatocyte markers are highlighted in blue. Cholangiocyte markers are highlighted in green. (G) Gene essentiality scores for *SMARCA4* (left) and *SMARCA2* (right) in *KEAP1* (WT) and *KEAP1* (Mutant) samples from TCGA_DEPMAP_.

## Discussion

Hepatocytes are an inherently plastic cell type with the capacity to undergo cell fate transition^45^. For example, hepatocytes have been shown to undergo transdifferentiation to cholangiocytes in response to liver injury, during liver regeneration, and in the context of tumour initiation^8,46–51^. Major epigenetic, transcriptional, and metabolic reprogramming events have been associated with transdifferentiation, however the precise mechanisms involved are poorly understood. In this study, we demonstrate that hepatocyte-specific activation of NRF2 initiates an evolutionarily conserved process of hepatocyte to cholangiocyte transdifferentiation in a manner dependent on SWI/SNF chromatin remodelling complexes.

Given that the NRF2 pathway can be activated in response to a variety of cell-intrinsic (e.g. mutations) and cell-extrinsic (e.g. oxidative stress) factors, we propose that NRF2 is an underappreciated regulator of hepatocyte plasticity. Importantly, our studies show that NRF2-dependent changes in liver cell fate are reversible. The transient nature of liver cell plasticity in this context is reminiscent of changes in cell plasticity associated with recovery of tissue homeostasis during liver regeneration^52^. Given that NRF2 is widely recognised to play an important role in liver regeneration^53–57^, future studies examining the role of NRF2-driven changes in cell fate plasticity during regeneration are warranted. Moreover, the fact that NRF2-driven changes in liver cell fate plasticity are reversible opens the possibility that this process can be targeted to restore liver homeostasis.

Dysregulation of cell fate is a hallmark of liver tumour initiation^58^. Accumulating evidence in animal models suggests that CCA can arise from malignant hepatocytes transdifferentiating to cholangiocytes^3,6–9,59^. Previous work has reported that liver-specific compound loss of *KEAP1* and phosphatase and tensin homolog (PTEN) promotes cholangiocyte expansion^60^. Our study provides the first evidence that NRF2 pathway activation alone is sufficient to drive expansion of the cholangiocyte compartment. Moreover, our study reveals this process to be driven by transdifferentiation of hepatocytes to cholangiocytes, thereby identifying hepatocytes as the cell-of-origin for CCA in the context of constitutive NRF2 activation. Interestingly, bioinformatic analysis of whole genome sequencing and epigenetic features of clinical samples supports the idea that hepatocytes are the cell-of-origin for hepatitis-induced CCA^61^. In addition to CCA, other types of liver cancer, including combined hepatocellular-cholangiocarcinoma (CHC), have been shown to arise from transdifferentiated hepatocytes^62^. CHC is characterized by dysregulated liver cell fate exhibiting features of HCC and CCA in the same tumour^63^. Recently, NRF2 has been shown to contribute to the progression of the chronic liver disease metabolic dysfunction-associated steatohepatitis (MASH) to HCC via a disease-associated hepatocyte state characterised by loss of cell identity^15,64,65^.

Epigenetic remodelling plays a critical role in regulating cell fate^35–38^. It is therefore not surprising that disruption of epigenetic machinery plays an important role in tumorigenesis^66^. Components of SWI/SNF chromatin remodelling complexes constitute some of the most frequently altered epigenetic regulators in cancer^38,39^. In this study, we have shown that NRF2-driven changes in liver cell fate plasticity are mediated by SWI/SNF complexes, underscoring the importance of epigenetic alterations in liver cancer initiation. These insights were derived from experiments employing an epigenetic-focused chemical screen that identified the BRG1/BRM inhibitor FHD-286. Preclinical studies have highlighted the therapeutic potential of FHD-286 for the treatment of acute myeloid leukaemia (AML), where it promotes differentiation of leukaemic stem cells, and small cell lung cancer (SCLC) driven by POU domain, class 2, transcription factor 3 (POUF2F3)^67,68^. Studies revealing that NRF2-dependent induction of enhancer of zeste homolog 2 (EZH2) or FACT complex subunit 16 (SUPT16H) accelerates tumour initiation and progression underscore the importance of epigenetic remodelling to the oncogenic functions of NRF2^20,69^. Given that epigenetic remodelling involves the coordinated activities of multiple epigenetic complexes, it would be important in future studies to explore the role of other epigenetic regulators identified from our screen.

In conclusion, this study has revealed that NRF2 drives cell fate plasticity via mechanisms involving epigenetic remodelling during liver tumour initiation. Given that the NRF2 pathway is frequently mutated in other solid tumours, including lung cancer, it would be interesting to explore a broader role for NRF2 in mediating cell fate plasticity in other tissues.

## Materials and Methods

### Cell culture

HepG2 cells were purchased from CellBank Australia and maintained in MEM, NEAA, no glutamine (ThermoFisher Scientific, 10370) supplemented with 1 mM sodium pyruvate, 2 mM glutamine, and 10% foetal bovine serum (FBS). 293T cells were acquired from American Type Culture Collection (ATCC) and maintained in DMEM, high glucose (ThermoFisher Scientific, 11965) supplemented with 1 mM sodium pyruvate and 10% FBS. Cells were maintained in a humidified incubator at 37°C with 5% CO_2_. Cell lines were routinely assayed for mycoplasma contamination.

### Plasmids

EGFP and FLAG/HA-tagged-NRF2^T80K^ were cloned into the pLIX_403 plasmid (a gift from David Root, Addgene plasmid #41395) via Gateway Cloning to generate a lentiviral doxycycline-inducible expression vector. Guide RNAs for *KEAP1*, or a control targeting *AAVS1*, were cloned into the LentiCRISPRv2 vector (a gift from Feng Zhang, Addgene plasmid #52961) as previously described^24^. Lentiviral particles were generated by co-transfecting 293T cells with the aforementioned vectors and the third-generation lentiviral packaging vectors pMDLg/pRRE, pRSV-Rev, and pMD2.G (gifts from Didier Trono, Addgene plasmids #12251, #12253 and #12259). HepG2 cells were transduced with lentiviral particles for 48 hrs prior to selection with 3 μg/mL puromycin.

### Zebrafish Husbandry

Zebrafish were maintained according to institutional Animal Experimentation and Ethics Committee (AEC) guidelines (Approval E580, E666, E634, AEC2024-03). The Tg(*gstp1*:EGFP) line has been described in previous studies^70^. Transgenic lines generated in this study: Tg(*fabp10a*:rtTA-T2A-CFP-CAAX;*TRE*:NRF2^T80K^-P2A-NLS-mCherry)^uom308^, referred to as TO:NRF2^T80K^, and TgKI(mScarlet3-P2A-anxa4)^uom309^, referred to as *anxa4*:mScarlet3. Generation of compound *keap1a*/*keap1b* crispants, via CRISPR/Cas9 gene editing, was previously described^24^. For all experiments, clutch siblings were used as controls and all embryos and larvae were maintained at 28°C throughout development. Male zebrafish were used for adult experiments while sex of the organism was not determined for experiments performed at the larval stage.

### Generation of TgKI(mScarlet3-P2A-anxa4)

An Alt-R CRISPR-Cas9 crRNA (Integrated DNA Technologies) targeting *anxa4* (5’-CTTCATCTCTACAATGGCAG-3’) was complexed with recombinant Alt-R® S.p. Cas9 Nuclease V3 (Integrated DNA Technologies). The donor dsDNA for mScarlet3-p2a knock-in was amplified from pMCS-mScarlet3-p2a-HygroR (VectorBuilder) (Forward primer: 5’-C*T*CCAGACTGAAACCTTTTCTTCATCTCTACAATGgatagcaccgaggcagtgat-3’; Reverse primer: 5’-T*A*AAAATCTTCTCTTTGTTATTTCACTTACCGCTGCaggcccggggttttcttcaa-3’). Primers (Integrated DNA Technologies) with 5’ AmC6 modification (denoted by *) included a 35-bp homology sequence flanking the *anxa4* target site and a 20-bp homology sequence to mScarlet3-p2a (Fig. S1D). 300 ng of amplified dsDNA was added to Cas9-guide ribonucleoprotein complex to make a 5 µL injection mix. The mix (2 nL) was injected into one-cell stage zebrafish embryos.

### Doxycycline treatment

For cell-based studies, HepG2 cells were treated with media containing 200 ng/mL of doxycycline (DOX; Sigma, D5207). Media was replaced every 48 hrs. For larval studies, a maximum of 50 larvae were maintained in petri dishes containing 40 mL of E3 media and 10 mL of freshly harvested paramecia (*Paramecium caudatum*, 10 cm dishes from 5 to 10 dpf). Larvae were treated with either 0.1% DMSO or 7.5 µg/mL DOX in E3 media from 5 dpf. Media was replaced every 48 hrs. For adult studies, zebrafish were placed in 1 L breeder boxes and treated with either 0.1% DMSO or 7.5 µg/mL DOX in water for the indicated amount of time. Water was replaced every 24 hrs and feeding was performed daily.

### Histology

Zebrafish larvae or adult liver tissue were fixed in 4% paraformaldehyde (PFA) overnight at 4°C. Zebrafish larvae were embedded in agarose larval arrays after fixation. Samples were subjected to paraffin-embedding and serial sectioning. Sections were stained with hematoxylin and Eosin (H&E) or Masson’s trichome by the Peter MacCallum Cancer Center (PMCC) Centre for Advanced Histology and Microscopy (CAHM) according to standard protocols. Sections were imaged with an Olympus BX53 microscope.

### Periodic Acid-Schiff (PAS) staining

Deparaffinized and hydrated sections were stained using a PAS staining kit (Abcam, ab150680) according to manufacturer’s instructions. Briefly, sections were immersed in periodic acid solution for 10 mins followed by four washes with distilled water. Sections were then treated with Schiff’s solution for 30 mins followed by rinsing with hot running tap water. Finally, Light Green solution was applied for 2 mins as a counterstain. Slides were rinsed in distilled water and dehydrated before mounting. Sections were imaged with an Olympus BX53 microscope.

### Terminal deoxynucleotidyl transferase biotin–dUTP nick end labelling (TUNEL) assay

Deparaffinized and hydrated sections were stained using the *In situ* Cell Death Detection Kit, TMR red (Sigma-Aldrich, 12156792910) according to manufacturer’s instructions. Briefly, sections were permeabilized with 0.1% Triton X-100 in PBS for 20 mins followed by two washes with PBS. The section used as a positive control was treated with DNase I (Zymo Research, E1010) at room temperature for 10 mins. Sections were then washed with PBS, treated with TUNEL reaction mixture, and incubated in the dark for 60 mins at 37°C. Sections were mounted with a coverslip and imaged using an Olympus FV3000 microscope.

### Immunofluorescent staining of sections

For immunofluorescence analysis, sections were initially unmasked using a citrate buffer antigen protocol, as previously described^71,72^. Slides were subsequently stained with primary antibodies against GFP (Abcam, ab13970; 1:400), and 2F11 (anti-Anxa4; Abcam, ab71286; 1:300). Slides were incubated overnight at 4°C with primary antibodies diluted in PBS containing 5% Bovine Serum Albumin (BSA; Sigma Aldrich, A7906). Slides were subsequently washed three times with PBS containing 0.1% Tween (PBST). Antibody binding was visualised with secondary antibodies from LI-COR Biosciences: Goat Anti-Mouse IgG Polyclonal Antibody (IRDye® 680RD) and Goat Anti-Rabbit IgG Polyclonal Antibody (IRDye® 800CW). Sections were imaged with an Olympus FV4000 confocal microscope equipped with Olympus FV30-SW software.

### RNA extraction and Bulk RNA-Seq analysis

RNA was extracted from HepG2 cells using the NucleoSpin RNA kit (Macherey-Nagel, 740955) as per manufacturer’s instructions. Dissected liver tissues were flash frozen in liquid nitrogen and homogenized in TRIzol™ (Thermo Fisher Scientific, 15596018) before RNA was extracted using the Direct-zol™ RNA MiniPrep Kit (Zymo Research, R2051) according to manufacturer’s instructions. In all cases, RNA quality was confirmed using an Agilent 4200 TapeStation System. Libraries were prepared using the QuantSeq 3ʹ mRNA-Seq kit (Lexogen) and sequenced with an Illumina NextSeq 500, with single-end 100bp reads to a depth of 5M reads per sample. FASTQ files were uploaded to the Galaxy web platform for quality control (FastQC), trimming (Cutadapt), alignment (RNA STAR), and counting (featureCounts)^73^. Reads from HepG2 cells were aligned to the human genome assembly (Ensembl hg19, GRCh37). For zebrafish samples, reads were aligned to the zebrafish reference genome (Ensembl danRer11, GRCz11) and annotated with the GTF from Lawson *et. al.*^74^. Analysis of differentially expressed genes was performed with Limma-Voom (v3.40.6). Gene set enrichment analysis (GSEA) was performed and plotted in R 10 using the ‘clusterprofiler’ package^75^. Heatmaps and volcano plots were generated in R 10 using the ‘pheatmap’ and ‘ggplot’ packages respectively.

### Confocal imaging of zebrafish larvae

Zebrafish larvae were anesthetized in E3 medium containing 0.2 mg/mL tricaine and embedded in 1% low-melt agarose on 35 mm glass bottom dishes (MatTek, P35GTOP020C). Live imaging was performed on a Nikon SoRa Spinning Disk confocal microscope using a 40x water immersion objective equipped with the NIS Elements software. 3D rendering of the bile duct network was performed using the ‘Surface’ function on Imaris v9.3 (Oxford Instruments).

### In vivo epigenetic-focused chemical screen

TO:NRF2^T80K^; *anxa4*:mScarlet3 transgenic larvae were treated with either DMSO, 15 µg/mL DOX, or 15 µg/mL DOX in the presence of 1 µM of epigenetic inhibitor from 5 to 9 dpf. Larvae were maintained in E3 medium and fed paramecia throughout the experiment. Media was replaced every 48 hrs. At 9 dpf, larvae were fixed in 4% PFA at 4°C in the dark overnight. Larvae were then washed with PBST and imaged as described in the ‘*Confocal imaging of zebrafish larvae’* section. Images were analysed and quantified as described in the ‘*Image Analysis and Quantification’* section.

### Image Analysis and Quantification

Images acquired by confocal microscopy of tissue sections (adult liver) or maximum intensity projections (larval liver) were pixel-classified using Ilastik software^76^ to generate prediction masks for Anxa4 (cholangiocyte) and CFP (total liver) areas. The prediction masks were then processed via Cellprofiler^77^ to quantify cholangiocyte area (the ratio of Anxa4 area to total CFP area).

### Cell lysis and immunoblotting

To prepare whole cell lysates, HepG2 cells were washed with PBS and lysed in SDS lysis buffer (1% SDS, 50 mM Tris-HCl, 10 mM EDTA) containing a protease inhibitor cocktail (Sigma-Aldrich, P8340) and Pierce^TM^ Universal Nuclease (ThermoFisher Scientific, 88702). To prepare nuclear fractions, HepG2 cells were trypsinized and resuspended in Buffer A (pH 7.0, 10 mM HEPES, 5 mM MgCl2, 25 mM KCl) at a density of 1 × 10^4^ cells/μL. The cell suspension was subsequently passed twenty times through a 26-gauge needle and incubated on ice for 15 mins. NP-40 was added to each sample to attain a final concentration of 0.2%. After vortexing, lysates were centrifuged at 14,000 *g* for 1 min at 4°C. The resulting supernatant was removed, while the nuclear pellet was lysed in 1 × Laemmli buffer containing 5% β-mercaptoethanol and boiled at 95°C for 12 mins. Whole cell or nuclear lysates were resolved by SDS-PAGE and transferred to nitrocellulose membrane as previously described^24^. Membranes were incubated with anti-HNF4α (C11F12) Rabbit mAb (Cell Signaling Technology, 3113), anti-β-Actin (8H10D10) Mouse mAb (Cell Signaling Technology, 3700), anti-NRF2 (E5F1A) Rabbit mAb (Cell Signaling Technology, 20733), or anti-Histone H3 Rabbit pAb (Abcam, ab1791) primary antibodies prior to incubation with IRDye secondary antibodies (LI-COR) and imaged using the Odyssey DLx imaging system (LI-COR).

### Single-cell RNA-seq (scRNA-Seq)

Adult zebrafish were treated with either DMSO or 7.5 µg/μL DOX for 3 weeks before being euthanized in E3 medium containing 4 mg/mL tricaine prior to submerging in ice-cold water for 1 min. For each treatment group, the superior lobes from two zebrafish were washed in PBS and transferred to an RNAse-free tube containing 750 μL of cell dissociation buffer (10 mg/mL collagenase in TESCA buffer) pre-heated to 32°C. Samples were incubated at 32°C for 20 min with constant mixing on a Thermo-Shaker (500 rpm; Thermo Fisher Scientific). Reactions were quenched by addition of 750 μL of 10% FBS in PBS. Dissociated cells were pelleted by centrifugation at 400 *g* for 5 mins at 16°C. Supernatant was discarded and cell pellets were resuspended in 1% FBS in PBS containing SYTOX™ Red dead cell stain (Invitrogen, S34859; 1:1000) and filtered through a FACS tube strainer on ice. Cell sorting was performed with a BD FACSAria™ Fusion 3 Sorter (BD Biosciences). 370,000 live events per treatment group were collected in Eppendorf tubes containing 100 μL of scRNA-Seq buffer (0.4% FBS in DPBS containing RNAse inhibitor). Sample preparation, gel-in-bead emulsion (GEM) generation, and library construction were performed according to the Chromium™ Single Cell 3’ user guide (10x Genomics) by the PMCC Molecular Genomics Core (MGC). Samples were then sequenced on a NovaSeq 6000 with a depth of 500 M reads per treatment group. Single cell data was processed using the 10x Genomics Cell Ranger software mapping to the GRCz11 zebrafish genome. Ribosomal and haemoglobin genes were removed. Quality control was performed using Seurat in R^78^. Cells with fewer than 500 unique genes and a mitochondrial RNA content greater than 50% were determined as low-quality cells and removed from subsequent analysis. Ambient RNA was removed using the SoupX^79^ package. Doublet cells were identified and removed using the package scDblFinder^80^ in R.

### Quantitative real time polymerase chain reaction

Complementary DNA (cDNA) was synthesized from 1 μg of RNA using an Applied Biosystems High-Capacity cDNA Reverse Transcription Kit (Thermo Fisher Scientific, 4368814) as per manufacturer’s instructions. The cDNA was diluted 1:20 prior to use. Quantitative real-time polymerase chain reaction (qPCR) was performed using Applied Biosystems Fast SYBR™ Green Master Mix (Thermo Fisher Scientific, 4309155) and an Applied StepOnePlus Real-Time PCR System (Thermo Fisher Scientific). Quantification of gene expression was analysed using the ^ΔΔ^Ct method^81^ normalized to *RPLP0*. Primers used in this study are outlined in Table 1.

**Table 1.**
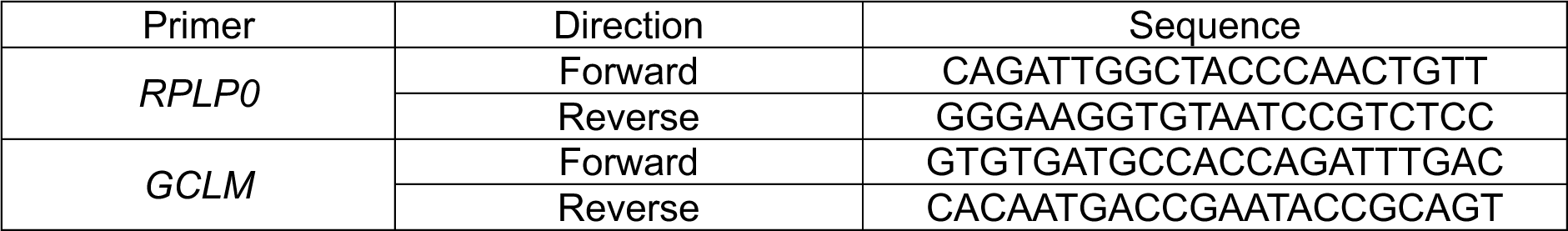
Primers used in the study.

### Gene essentiality analysis

The gene essentiality of *SMARCA2* and *SMARCA4* in *KEAP1* (WT) and *KEAP1* (mutant) cancers within TCGA_DEPMAP_ were analysed in Prism 10 software (GraphPad Software) based on data obtained from the Translational Dependency Tool (https://xushiabbvie.shinyapps.io/TDtool/)^44^.

### Statistical analyses

Statistical analyses were performed with Prism 10 software (GraphPad Software). All statistical analyses for data comparing two groups were performed with an unpaired Student’s t-test. One-way ANOVA with the Holm–Sidak method for multiple comparisons was used for comparison of more than two groups. All immunoblots are representative of results from at least three independent experiments. All other statistical details can be found in the figure legends.

## Acknowledgements

A.J.S.O. is supported by a Peter MacCallum Cancer Centre Foundation Grant. K.K.B. is supported by NHMRC Ideas Grants (GNT2004212 and GNT2012313) and a Victorian Cancer Agency Mid-Career Research Fellowship (MCRF17020). A.G.C. is supported by a National Health and Medical Research Council (NHMRC) Investigator Grant (GNT1176650), an Australian Research Council Discovery Project Grant (DP200102693), a GESA Project Grant, a Dame Kate Campbell Fellowship and a Victorian Cancer Agency Mid-Career Research Fellowship (MCRF23013). K.K.B and A.G.C. are also jointly supported by the Peter MacCallum Cancer Foundation (Ted and Lila Seehusen Foundation). We acknowledge support from the Peter MacCallum Cancer Centre Foundation and the Australian Cancer Research Foundation. We thank the staff at the University of Melbourne Zebrafish Core Facility for their support and contributions. We also extend our thanks to the Peter MacCallum Cancer Centre Core Facilities and their staff who provided support for this work; namely the Molecular Genomics Core (RRID:SCR_025695), the Centre for Advanced Histology and Microscopy (RRID:SCR_025432), and Research Laboratory Support Services (RRID:SCR_025699). Finally, we thank members of the Cox Laboratory and Brown Laboratory (Peter MacCallum Cancer Centre) for helpful discussions.

## Author contributions

A.J.S.O., A.G.C., and K.K.B. designed research; A.J.S.O. and A.P.K. performed research; A.J.S.O. and K.J.E. analyzed data; A.G.C. and K.K.B. provided supervision; and A.J.S.O., A.G.C., and K.K.B. wrote the paper.

## Supplementary Figures

**Figure S1.**
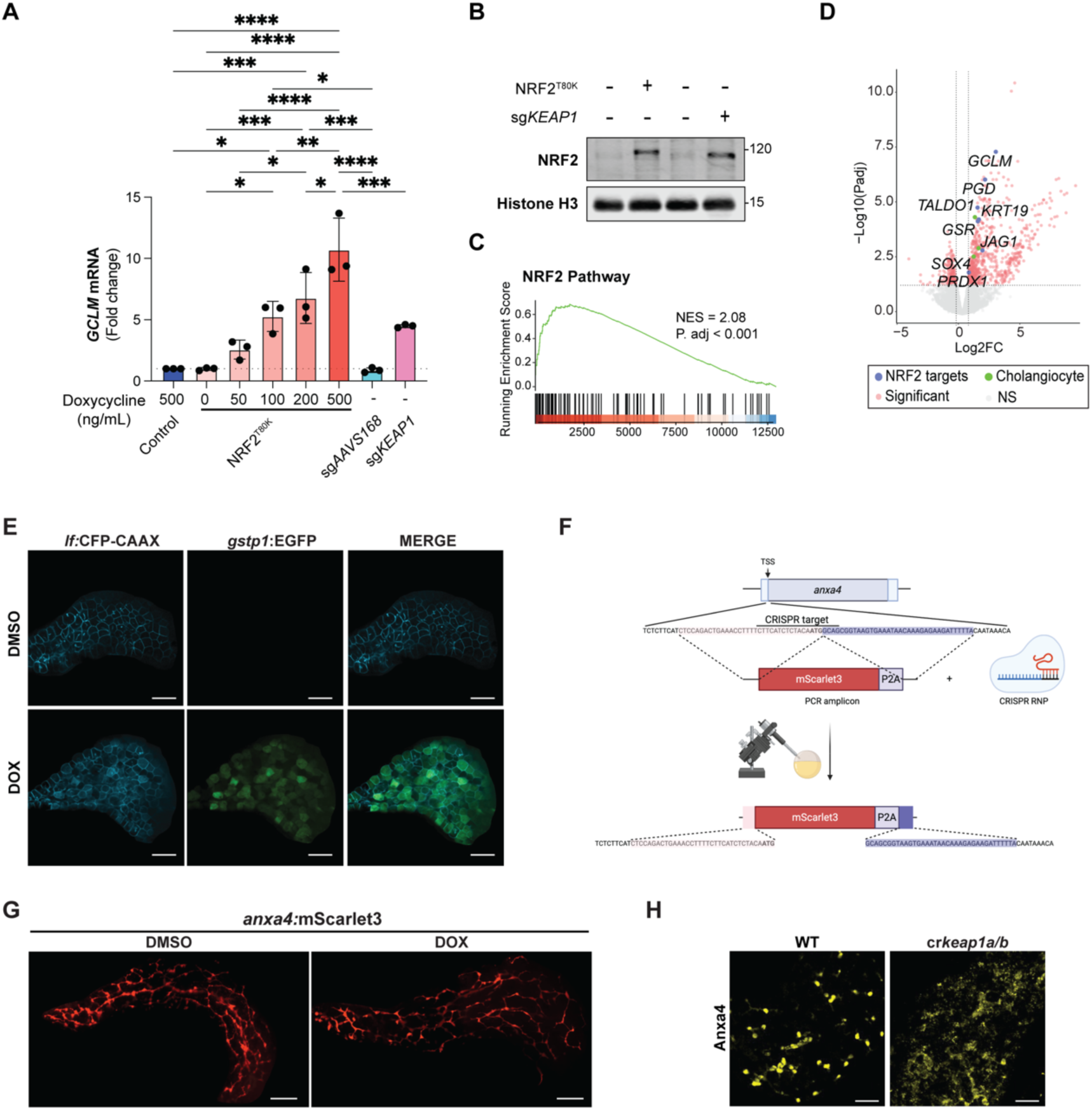
Constitutive NRF2 activation promotes liver cell plasticity *in vitro* and *in vivo*. (A) Quantitative PCR (qPCR) analysis of *GCLM* expression in Control and TO:NRF2^T80K^ HepG2 cells treated with DOX for 4 days, and sgAAVS1 and sg*KEAP1* HepG2 cells, n=3. (B) Representative immunoblot analysis of nuclear fractions from Control and TO:NRF2^T80K^ HepG2 cells treated with 200 ng/mL DOX for 4 days and sgAAVS1 and sg*KEAP1* HepG2 cells. (C) Representative confocal images of CFP (hepatocyte, cyan) and EGFP (NRF2 target gene expression, green) from TO:NRF2^T80K^;*gstp1:*EGFP zebrafish larvae treated with DMSO or DOX from 5 to 6 dpf. Scale bars represent 50 µm. (D) Schematic of the approach employed to generate the TgKI(mScarlet3-P2A-anxa4) transgenic zebrafish line. (E) Representative confocal images of mScarlet3 (cholangiocyte) from *anxa4*:mScarlet3 zebrafish larvae at treated with DMSO or DOX from 5 to 10 dpf. Scale bars represent 50 µm. (F) Immunofluorescent staining of Anxa4 (cholangiocyte) in liver sections from WT and *keap1a/b* crispants at 14 dpf. Scale bars represent 25 µm.

**Figure S2.**
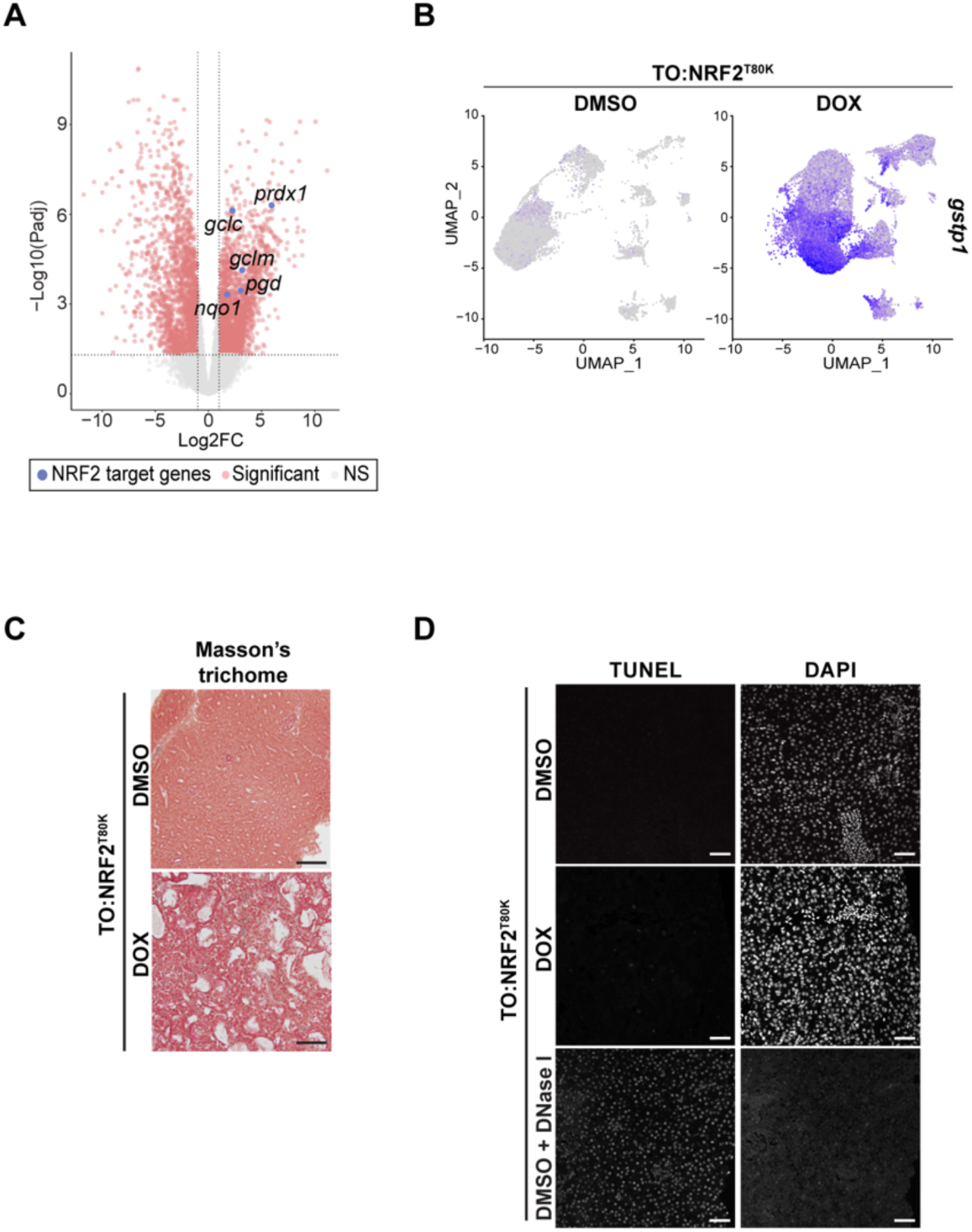
NRF2 promotes transdifferentiation of hepatocytes to cholangiocytes. (A) Volcano plot of DEGs identified by bulk RNA-Seq analysis of livers from adult TO:NRF2^T80K^ zebrafish treated with DOX versus DMSO for 4 weeks, n=3. Significant DEGs are highlighted in pink. Select canonical NRF2 target genes are highlighted in blue. (B) UMAP visualization of *gstp1* expression in livers from adult TO:NRF2^T80K^ zebrafish treated with DMSO or DOX for 3 weeks. (C) Masson’s trichome stained histological sections from livers isolated from adult TO:NRF2^T80K^ zebrafish after 4 weeks treatment with DMSO or DOX. Scale bars represent 50 µm. (D) TUNEL- and DAPI-stained histological sections from livers isolated from adult TO:NRF2^T80K^ zebrafish after 4 weeks treatment with DMSO or DOX. DNase I treatment was employed as a positive control. Scale bars represent 50 µm.

**Figure S3.**
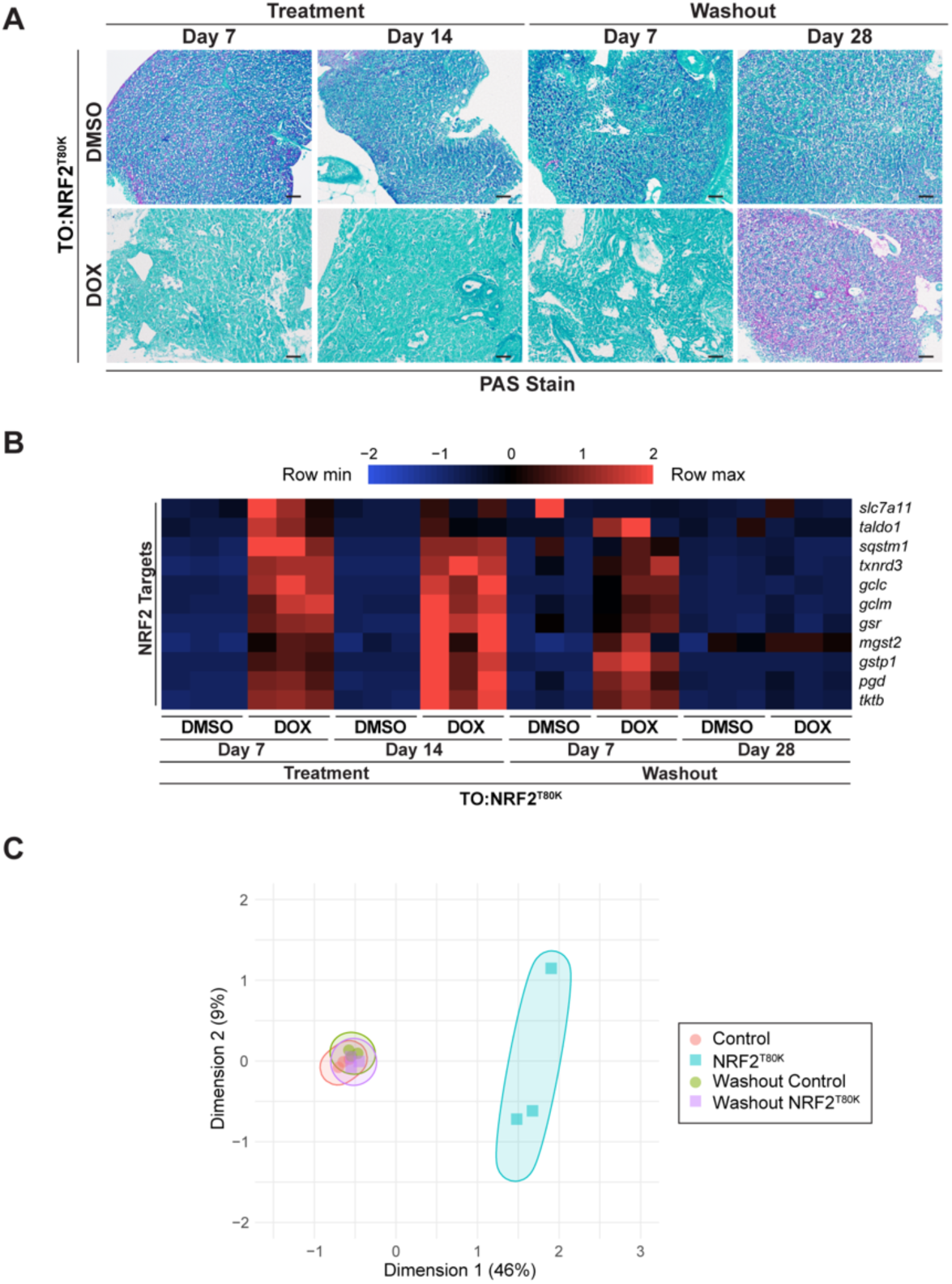
NRF2 is required to maintain altered liver cell fate identity. (A) Periodic Acid-Schiff (PAS)-stained histological sections from livers isolated from adult TO:NRF2^T80K^ zebrafish after 7- or 14-days treatment with DMSO or DOX, followed by 7- or 28-days withdrawal of DMSO or DOX. Scale bars represent 50 µm. (B) Heatmap of NRF2 target gene expression in livers isolated from adult TO:NRF2^T80K^ zebrafish after 7- or 14-days treatment with DMSO or DOX, followed by 7- or 28-days withdrawal of DMSO or DOX as determined by bulk RNA-Seq analysis, n=3. (C) MDS Plot of RNA-Seq data from Control and TO:NRF2^T80K^ HepG2 cells after 4 days of DOX treatment, followed by withdrawal of DOX for 4 days.

**Figure S4.**
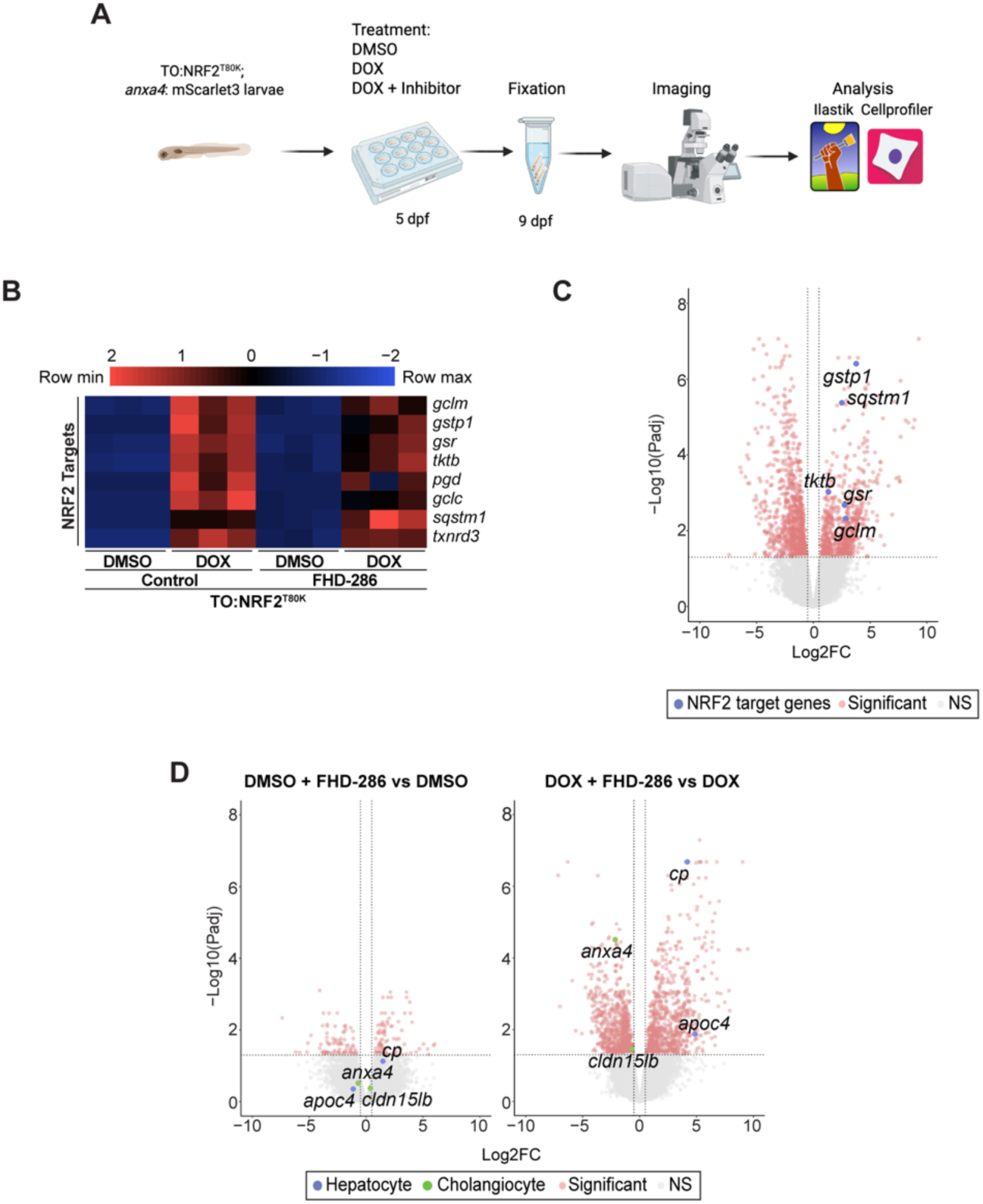
Epigenetic-focused small molecule inhibitor screen reveals that NRF2-driven liver cell plasticity is mediated by BRG1/BRM. (A) Schematic of the epigenetic-focused small molecule inhibitor screen. (B) Heatmap of NRF2 target gene expression in livers isolated from adult TO:NRF2^T80K^ zebrafish treated with DMSO or DOX in the absence or presence of 250 nM FHD-286 for 14 days as determined by bulk RNA-Seq analysis, n=3. (C) Volcano plot of DEGs identified by bulk RNA-Seq of livers isolated from adult TO:NRF2^T80K^ zebrafish treated with DOX versus DMSO both in the presence of 250 nM FHD-286 for 14 days as determined by bulk RNA-Seq analysis, n=3. Significant DEGs are highlighted in pink. Select canonical NRF2 target genes are highlighted in blue.

**Supplementary Table 1.**
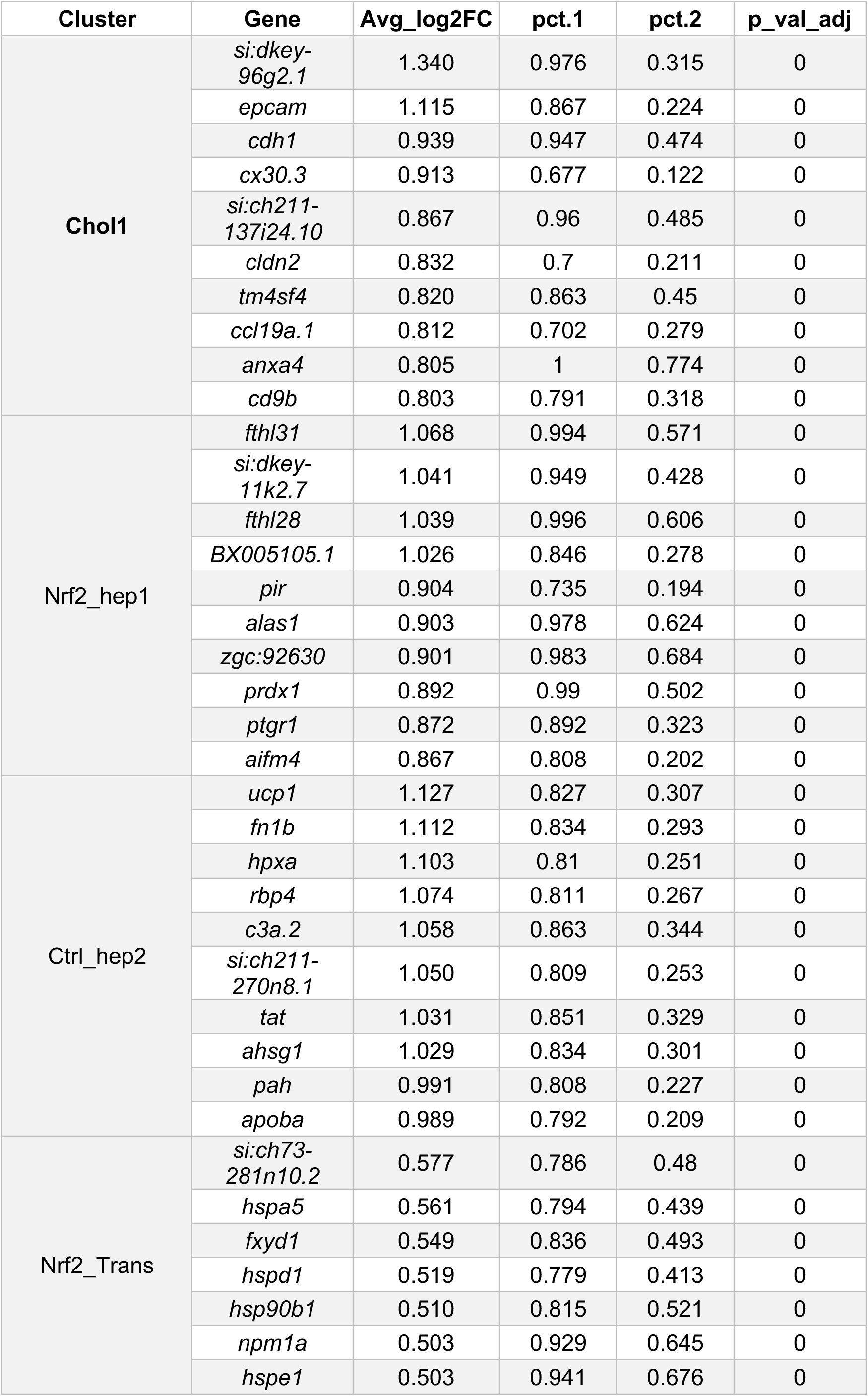
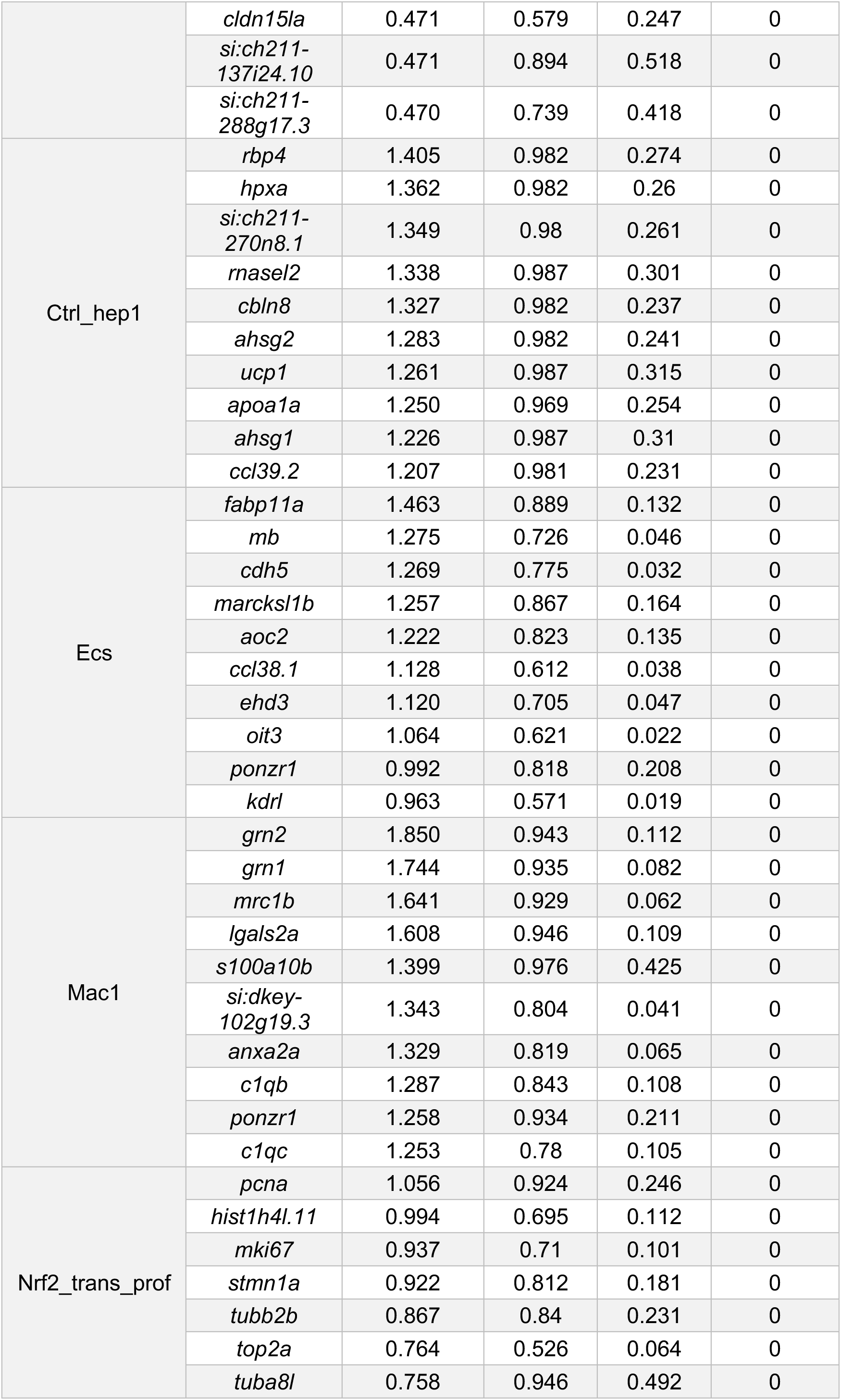
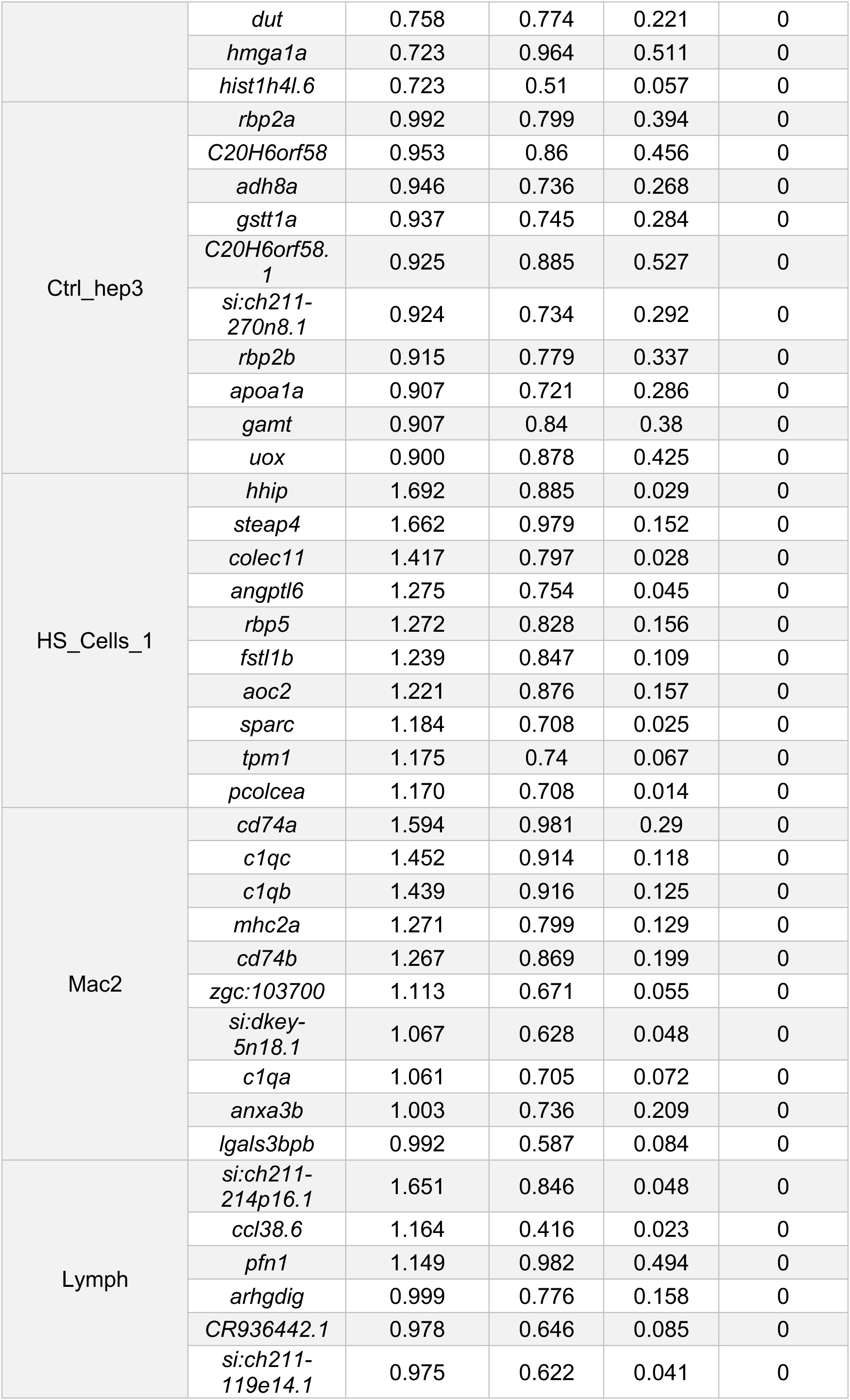
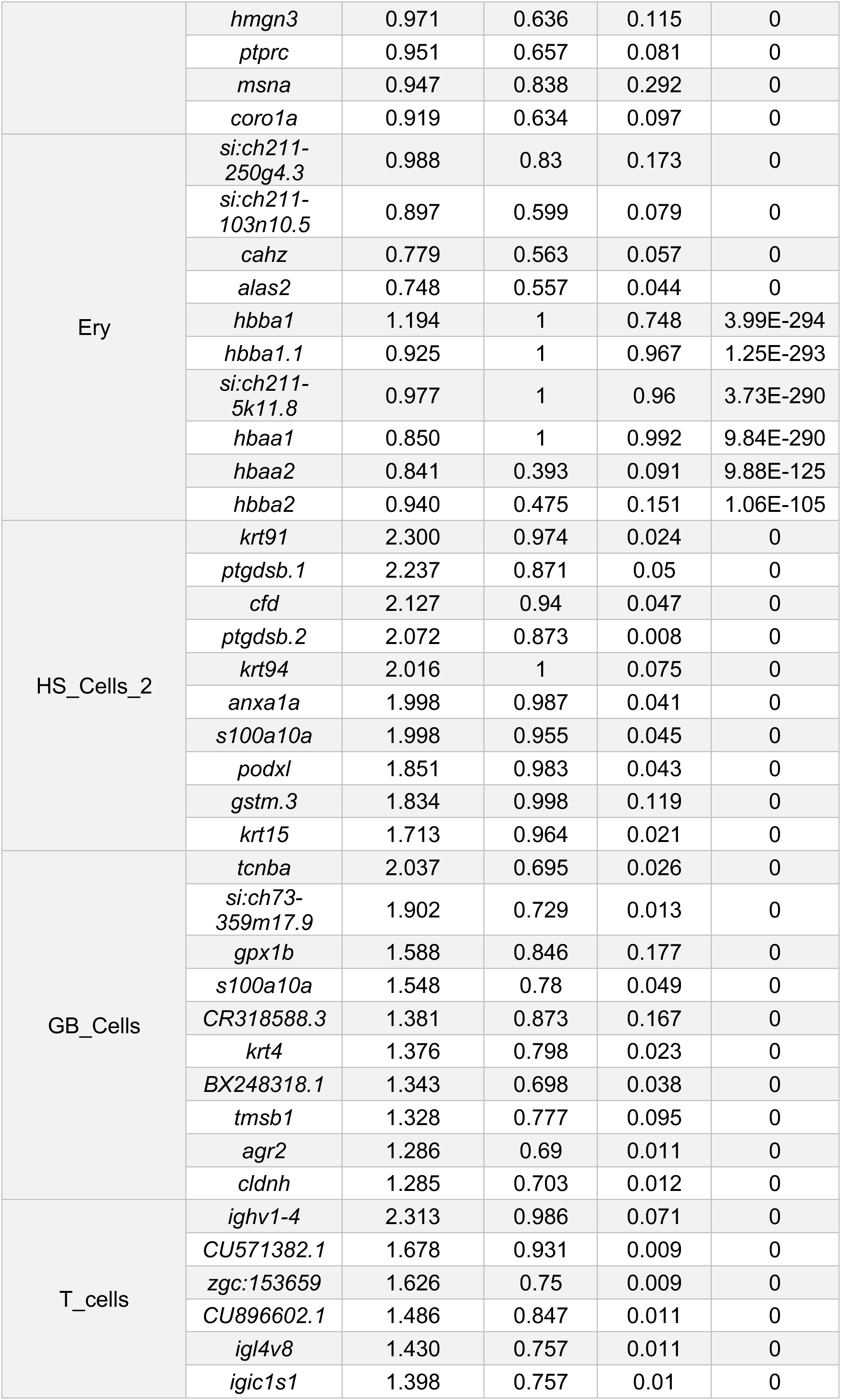
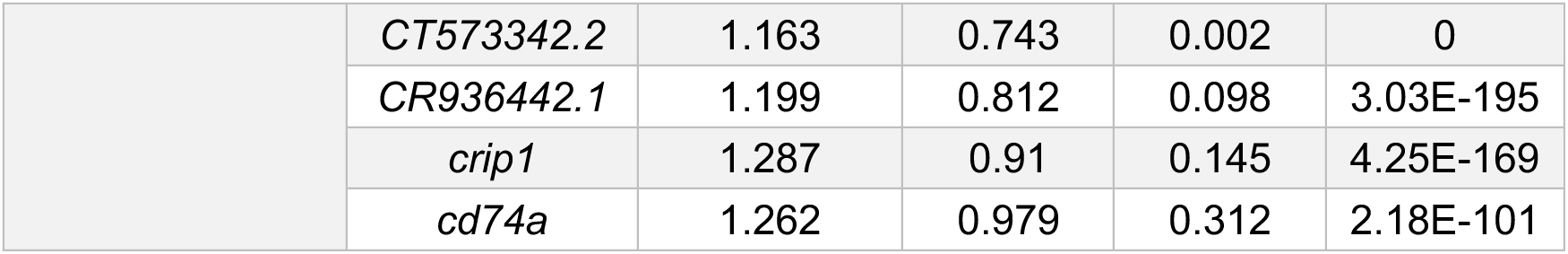
Top 10 differentially expressed genes in scRNA-Seq clusters.

**Supplementary Table 2.**
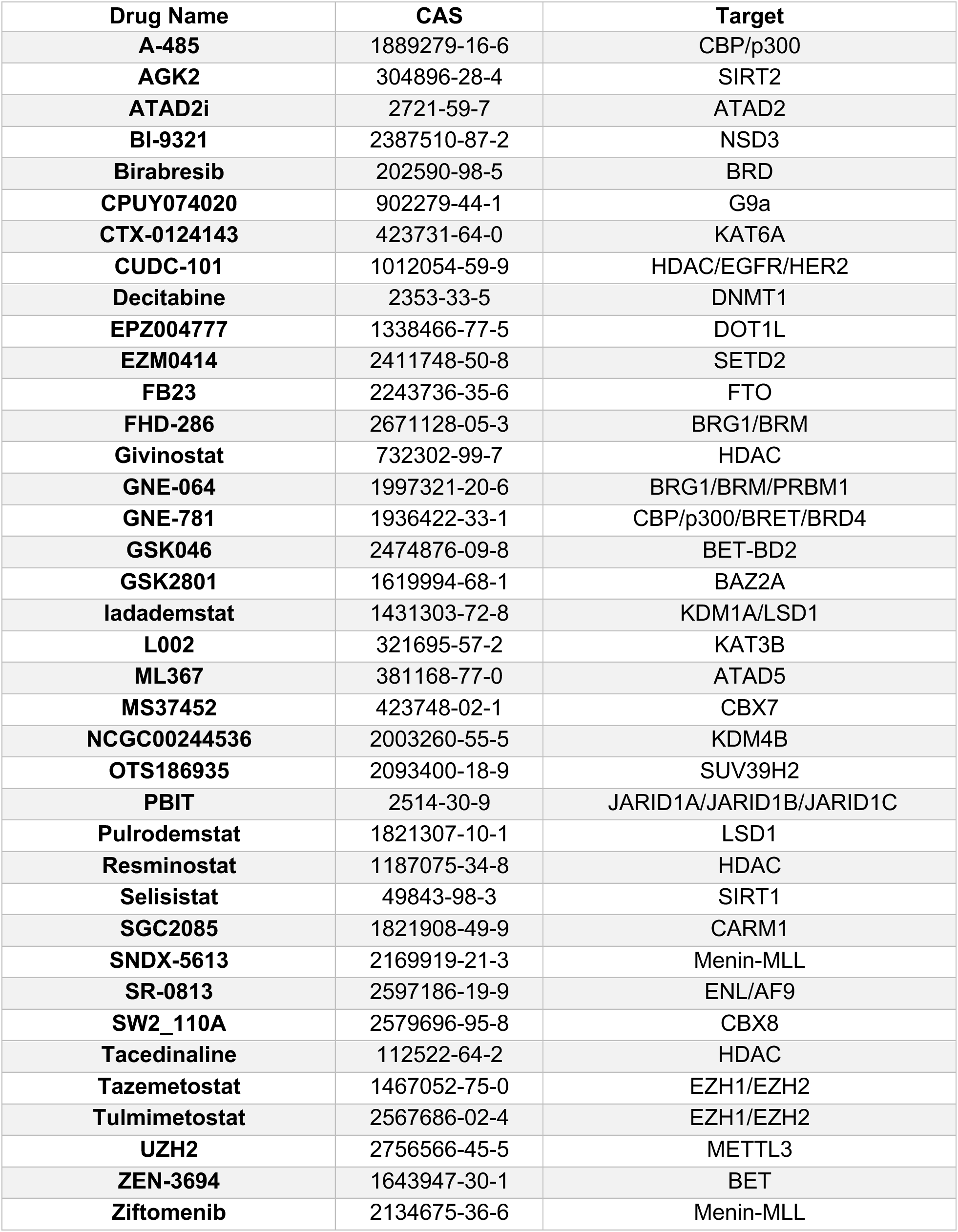
Small molecules in the epigenetic-focused chemical library.

